# Ecology, Not Distance, Explains Community Composition in Parasites of Sky-Island Audubon’s Warblers

**DOI:** 10.1101/346627

**Authors:** Jessie L. Williamson, Cole J. Wolf, Lisa N. Barrow, Matthew J. Baumann, Spencer C. Galen, C. Jonathan Schmitt, Donna C. Schmitt, Ara S. Winter, Christopher C. Witt

**Affiliations:** Museum of Southwestern Biology, University of New Mexico, 302 Yale Blvd NE, Albuquerque, NM 87131, USA; Department of Biology, University of New Mexico, MSC03-2020, Albuquerque, NM 87131, USA.; Division of Biological Sciences, University of Montana, 32 Campus Dr. HS 104, Missoula, MT 59812, USA.; Sackler Institute for Comparative Genomics, American Museum of Natural History, Central Park West at 79^th^ Street, New York, NY 10024, USA.; Department of Organismic and Evolutionary Biology and Museum of Comparative Zoology, Harvard University, 26 Oxford St., Cambridge, MA 02138, USA.

**Keywords:** beta diversity, community composition, *Leucocytozoon*, *Parahaemoproteus*, parasite biogeography, *Plasmodium*, *Setophaga auduboni*, species turnover

## Abstract

1. Haemosporidian parasites of birds are ubiquitous in terrestrial ecosystems, but their coevolutionary dynamics remain poorly understood. If species turnover in parasites occurs at a finer scale than species turnover in hosts, widespread hosts would encounter diverse parasites and potentially diversify as a result. Previous studies have shown that some wide-ranging hosts encounter varied haemosporidian communities throughout their range, and vice-versa. However, it remains difficult to test spatial patterns of diversity in this complex multi-host multi-parasite system because it remains inadequately surveyed.
2. We sought to understand how and why a community of avian haemosporidian parasites varies in abundance and composition across an array of eight sky islands in southwestern North America. We tested whether bird community composition, aspects of the environment, or geographic distance explain parasite species turnover in a widespread, generalist host.
3. We sampled 178 Audubon’s Warblers (*Setophaga auduboni*) along elevational transects in eight mountain ranges and screened them for haemosporidian mtDNA. We tested predictors of infection using generalized linear models (GLMs) and we tested predictors of bird- and parasite-community dissimilarity using generalized dissimilarity modeling (GDM).
4. Predictors of infection differed by genus: *Parahaemoproteus* was predicted by elevation and climate, *Leucocytozoon* varied idiosyncratically among mountain ranges, and *Plasmodium* was unpredictable, but rare. Parasite species turnover was nearly three-fold higher than bird species turnover and was predicted by elevation, climate, and bird community composition, but not by geographic distance.
5. Haemosporidian communities vary strikingly at spatial scales of hundreds of kilometers, across which the bird community varies only subtly. The finer spatial scale of turnover among parasites species implies that their ranges tend to be smaller than those of their hosts. Avian host species should encounter different parasite species in different parts of their ranges, resulting in spatially varying selection on host immune systems. Furthermore, the fact that parasite turnover was predicted by bird turnover implies that different species within a host community affect each other’s parasites, potentially facilitating indirect antagonistic effects.

## INTRODUCTION

Spatial variation among parasite communities provides a critical window into ecology and evolution of host-parasite dynamics. The degree to which parasite community composition is affected by climate, host community, geographic distance, or dispersal barriers remains poorly understood for most parasites. Community similarity generally declines with distance, referred to as the ‘distance-decay’ relationship (Nekola & White 1999; Poulin 2003), and it is thought that these relationships and their underlying predictors differ between micro-organisms, such as parasites, and macro-organisms, such as hosts (Astorga *et al.* 2012; Nemergut *et al.* 2013). Turnover, or change in community composition across space, might be closely linked for host and parasite communities because of the total dependence of parasites on hosts. This has been demonstrated previously for haemosporidian parasites of birds (Ishtiaq *et al.* 2010; Svensson-Coelho & Ricklefs 2011; Clark & Clegg 2017). However, there are several possible reasons to expect mismatches between host and parasite turnover.

Species traits such as dispersal ability or niche breadth should affect rates of turnover, and these traits could differ systematically between hosts and parasites. Parasite dispersal potential is often believed to be greater than that of hosts, because potential for dispersal at different life stages or in different host species is possible (Njabo *et al.* 2011; Carlson *et al.* 2015). Additionally, host-generalist parasites may have opportunities to expand their geographic ranges beyond individual host species, whose ranges are constrained by forces that may not affect the parasite (Poulin 2003). These mechanisms would predict lower parasite turnover relative to host turnover. On the other hand, there are several reasons why parasite species turnover could be higher than that of hosts: (1) Parasite dispersal may be reduced because they are sensitive to environmental clines, as when conditions for parasite transmission or reproduction (e.g. vectors) are limited spatially or seasonally (Njabo *et al.* 2011); (2) particular reproductive modes (e.g. longer times in the free-living stages, or shorter generation times) could facilitate faster diversification in parasites (Mazé-Guilmo *et al.* 2016); and (3) parasites can become physically isolated by host-switching within geographic regions, causing accumulation of parasite diversity at a smaller spatial scale than in hosts (Galen and Witt 2014).

Haemosporidians (Apicomplexa: Haemosporida) are a cosmopolitan clade of intracellular blood parasites that includes at least three genera that infect songbirds: *Parahaemoproteus*, *Plasmodium*, and *Leucocytozoon* (Galen *et al.* 2018). These parasites can affect hosts by diminishing survival and condition (Marzal *et al.* 2008; Asghar *et al.* 2015), reducing reproductive success (Marzal *et al.* 2005; Knowles, Palinauskas & Sheldon 2010), and potentially accelerating diversification of allopatric or parapatric populations that are exposed to different parasite faunas (Ricklefs 2010; Thornhill & Fincher 2013). Haemosporidians are thought to diversify rapidly via host-switching (Ricklefs & Outlaw 2010; Galen & Witt 2014; Ricklefs *et al.* 2014), but the roles of geography and physical isolation in diversification remain poorly understood. Some haemosporidian parasites are closely associated with a single host species, while others appear to be broadly-distributed generalists (Moens & Pérez-Tris 2016; Soares, Latta & Ricklefs 2017). In at least some cases, the distributions of haemosporidians are thought to depend on avian host-species distributions (Reullier *et al.* 2006), vector abundance (Kimura, Dhondt & Lovette 2006), or climate (Sehgal *et al.* 2010). Although over 3,000 unique lineages of avian haemosporidians have been documented (Bensch, Hellgren & Pérez-Tris 2009), it is not yet possible to compare host and parasite richness because the vast majority of haemosporidian-host interactions remain undescribed. One repeated finding from previous studies in both tropical and temperate zones is that there are differences among the three haemosporidian genera in their biogeographic tendencies: *Plasmodium* lineages tend to be broadly-distributed host-generalists (e.g. Walther *et al.* 2016), *Parahaemoproteus* lineages tend to be more host-specialized (e.g. Moens & Pérez-Tris 2016), and *Leucocytozoon* lineages tend to be sensitive to climate (e.g. Galen & Witt 2014).

The spatial scale and potential drivers of haemosporidian turnover have been tested in both island and continental systems, with variable results (Svensson-Coelho & Ricklefs 2011; Scordato & Kardish 2014; Ellis *et al.* 2015; Fecchio *et al.* 2017b; Soares *et al.* 2017). Svensson-Coelho & Ricklefs (2011) found higher rates of turnover in Lesser Antillean parasite communities than bird communities, and that geographic distance explained parasite turnover in only one of three focal hosts, the geographically-restricted Black-faced Grassquit. Ishtiaq *et al.* (2010) found a distance-decay effect for *Plasmodium* but not *Parahaemoproteus* in Melanesia. Clark and Clegg (2017) found that host turnover explained parasite turnover in Melanesia, particularly in *Parahaemoproteus*, but also in *Plasmodium*. In eastern North America, Ellis *et al.* (2015) also found that host turnover affected parasite turnover, but that climate and geographic distance did not. Ellis et al. (2015) further found that haemosporidian lineages infecting Northern Cardinals varied from locality to locality more than expected based on turnover in the whole haemosporidian community, suggesting coevolution with different parasite lineages in different parts of the host range. Fecchio *et al.* (2017b) found that modest rates of turnover among haemosporidian communities in manakins (Pipridae) across the tropical Amazon were not predicted by distance, environment, or host turnover. In sum, previous studies show that the spatial scale of haemosporidian turnover and its causes remain poorly understood.

The sky islands of the arid southwestern United States are an ideal setting in which to test causes of turnover among parasite communities. The term ‘sky island’ refers to high-elevation forests that are geographically isolated by expanses of low, arid habitats (McCormack, Huang & Knowles 2009). Southwestern sky islands were most recently connected during the LGM (~18−9 Ka), although some sky-island clades diversified during earlier divergence events, >1 Mya (McCormack, Bowen & Smith 2008). It is not known whether parasite community variation among sky islands tracks host community variation or is governed by other factors.

Here we address patterns and drivers of haemosporidian turnover among sky islands separated by tens to hundreds of kilometers using generalized dissimilarity modeling (GDM). GDM, a form of nonlinear matrix regression, permits analysis of spatial patterns of community dissimilarity as a function of environmental dissimilarity and geographic distance (Ferrier *et al.* 2007; Fitzpatrick *et al.* 2013). It is an appropriate method to understand turnover because it accounts for expected non-linear relationships of community dissimilarity with environmental and geographic distance, respectively, and it is robust to collinearity (Fitzpatrick *et al.* 2013; Warren *et al.* 2014; Glassman, Wang & Bruns 2017). Using generalized linear modeling and GDM, we asked: (1) What, if anything, predicts avian haemosporidian infection in an array of southwestern sky islands? (2) Do climate, host community, and/or geographic distance explain turnover of haemosporidian lineages among these sky islands?

As a consistent and practical way of sampling sky-island haemosporidian communities, we surveyed parasites from a single, widespread and abundant host, the Audubon’s Warbler (*Setophaga auduboni; Aves:* Parulidae). This species breeds above ~2,100 m elevation in forests of southwestern North America, becoming more abundant at higher elevations (Hunt & Flaspohler 1998). *S. auduboni* exhibits little to no phenotypic variation (Milá, Smith & Wayne 2007) or population genomic differentiation (Toews *et al.* 2016) across the geographic extent of our study area. Given the tendency of haemosporidian parasites to infect multiple host species (Galen & Witt 2014; Moens *et al.* 2016), we expect that *S. auduboni* shares a large number of haemosporidian lineages with other hosts. A single, abundant host species is likely to provide a window into patterns and processes occurring in the whole haemosporidian community (Svensson-Coelho & Ricklefs 2011), as well as a direct test of spatial variation in parasitism experienced by a single host.

## METHODS

### Field sampling

We collected 178 *S. auduboni* specimens along elevational transects (~2,200−3,500 m) in eight sky island mountain ranges (Fig. 1, Table 1) in compliance with animal care regulations and federal, state, and tribal collecting permits. Specimens, along with frozen tissues and data, were deposited in the Museum of Southwestern Biology and are searchable via the open-access Arctos database (arctosdb. org; Table S1).

**Fig. 1.**
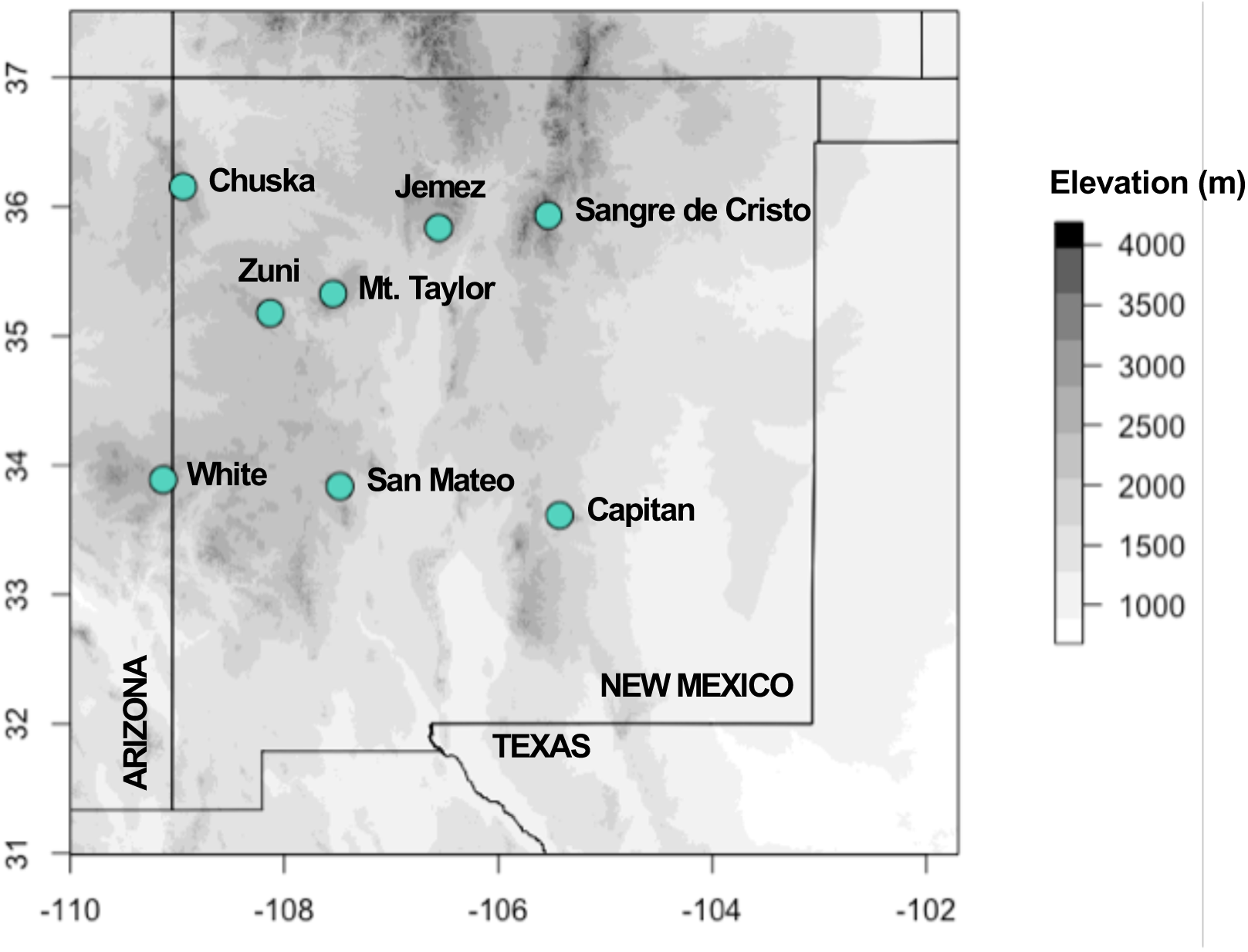
Map of sampled sky islands, with higher elevations shaded darker. Map data from Jarvis *et al.* (2008).

**Table 1.** Summary of sky island study site locations and elevational ranges. For each site we report the total number of: birds sampled, infections recovered, haplotypes recovered and haplogroups recovered.

| Site | Mean latitude and longitude | Elevational range of samples (m) | Birds sampled | Total infections | Number of Haplotypes | Number of Haplogroups |
| --- | --- | --- | --- | --- | --- | --- |
| Capitan | 33.614, -105.427 | 2286–3103 | 14 | 7 | 6 | 5 |
| Chuska | 36.156, -108.947 | 2223–2812 | 45 | 46 | 16 | 10 |
| Jemez | 35.837, -106.558 | 2489–3372 | 17 | 17 | 13 | 10 |
| Mt. Taylor | 35.3289, -107.546 | 2404–3321 | 26 | 32 | 16 | 13 |
| Sangre de Cristo | 35.933, -105.533 | 2283–3504 | 16 | 8 | 4 | 4 |
| San Mateo | 33.837, -107.481 | 2354–3064 | 18 | 15 | 9 | 7 |
| White | 33.888, -109.1329 | 2382–3060 | 25 | 16 | 9 | 8 |
| Zuni | 35.179, -108.132 | 2303–2766 | 17 | 17 | 9 | 8 |

### Genetic analysis

We extracted DNA from pectoral muscle with a QIAGEN kit. We used three nested PCRs to amplify a 478-bp fragment of mitochondrial *cytb* (Hellgren, Waldenström & Bensch 2004; Waldenström *et al.* 2004; see Appendix 1 in Supporting Information). Haplotypes differing by one or more base pairs (~0.2%) from published sequences on GenBank or MalAvi (Bensch *et al.* 2009) were considered novel (not found previously). Co-infections were phased with existing haplotypes when possible (Appendix S1). Novel haplotypes were named following MalAvi conventions, and sequences were uploaded to GenBank and MalAvi.

There is evidence that haemosporidian sequences that differ by as little as a single nucleotide may be different species (Bensch *et al.* 2009). Nonetheless, classifying lineages based on a one base pair cyt *b* difference could potentially lead to overestimates of parasite diversity. We analyzed all haplotypes as independent lineages and also adopted a haplotype-classification approach that allowed us to test our predictions under a more conservative criterion, modeled after (Svensson-coelho *et al.* 2013; Appendix S1). We collapsed all haplotypes into ‘haplogroups’, following a two-level scheme: (1) haplotypes were collapsed if they co-occurred in the same individual and differed by one base pair; and/or (2) haplotypes were collapsed if they overlapped in distribution (defined as co-occurrence in one or more sky islands) and differed by one base pair (Fig. S1; Tables S1 and S4). We conducted all analyses for both haplotypes and haplogroups.

### Modeling Haemosporidian Infection Status

We defined *infection status* as the total number of infected hosts, divided by the total number of screened hosts (proportion of individuals infected). We used Chi-square tests to evaluate effects of sex and age on infection status. We found no differences between sexes or ages; thus, we excluded age and sex from subsequent models.

We used principal component analysis (PCA) of WorldClim data v1.4 (Hijmans *et al.* 2005) to characterize climate variability among sky islands. The first two PCA axes (PC1 and PC2) explained 83.0% of climate variation and were subsequently used as predictor variables in models. Loadings indicated that PC1 represented increased temperature and decreased precipitation, which we refer to as “temperature-aridity index”. PC2 represented increased temperature seasonality and decreased precipitation, which we henceforth refer to as “seasonality index”.

To determine causes of infection status, we constructed a set of binomial GLMs for each haemosporidian genus. Each set included all possible additive combinations of five explanatory variables: site, elevation, latitude, temperature-aridity index (PC1), and seasonality index (PC2). We did not consider quadratic variables or interactions. Input variables were centered and scaled. Each model set included an intercept-only (null) model. We evaluated goodness-of-fit by examining residual deviance statistics for the global model in each set, and evaluated overdispersion using *ĉ*. We used AICc and Akaike weights (w*_i_*) to compare models. When *ĉ* was >1, quasi-AICc (QAICc) was used (Richards 2008). We eliminated models with uninformative parameters, defined as having a higher AICc or QAICc score than a nested version of the same model (Richards 2008; Arnold 2010; Table 2). The null model was retained in all candidate model sets, and the effect of each predictor was estimated by calculating model-averaged regression coefficients (Table 3). We tested 32 candidate models for each model set. Final models are presented in Table 2. All models were built in R, v3.3.2 (R Core Team 2016).

**Table 2.** Comparison of three sets of candidate models describing predictors of *Parahaemoproteus*, *Plasmodium*, and *Leucocytozoon* infection status. Models are ranked in ascending order by their ΔQAICc or ΔAICc scores relative to model with the lowest QAICc or AICc score in the set. *Parahaemoproteus* models were evaluated using QAICc and ΔQAICc; and *Plasmodium* and *Leucocytozoon* models using AICc and ΔAICc. Akaike weights (w_*i*_) quantify the probability that a particular model is the best model in the set, given the data. K indicates number of parameters. Model sets included models with single main effects and additive (+) effects of five explanatory variables: study site, elevation, latitude, temperature-aridity index (PC1), and seasonality index (PC2). Candidate sets were selected from full model sets by eliminating any model with a higher QAICc or AICc score than a nested version of the same model.

| Model Set | Model | K | QAICc or AICc | $\Delta\text{QAICc}$ or $\Delta\text{AICc}$ | $W_i$ |
| --- | --- | --- | --- | --- | --- |
| *Parahaemoproteus* | elevation + PC2 | 4 | 181.28 | 0.00 | 0.43 |
|  | PC1 + PC2 | 4 | 182.59 | 1.31 | 0.22 |
|  | PC1 | 3 | 183.57 | 2.29 | 0.14 |
|  | elevation + latitude | 4 | 183.86 | 2.58 | 0.12 |
|  | elevation | 3 | 185.61 | 4.33 | 0.05 |
|  | latitude | 3 | 187.94 | 6.66 | 0.02 |
|  | site + PC2 | 10 | 188.04 | 6.76 | 0.01 |
|  | PC2 | 3 | 189.63 | 8.35 | 0.01 |
|  | site | 9 | 190.27 | 8.99 | 0.00 |
|  | Intercept only | 2 | 190.94 | 9.66 | 0.00 |
| *Plasmodium* | Intercept only | 1 | 89.020 | 0.00 | 1.00 |
| *Leucocytozoon* | site | 8 | 154.83 | 0.00 | 0.98 |
|  | elevation | 2 | 164.68 | 9.85 | 0.01 |
|  | PC1 | 2 | 165.00 | 10.17 | 0.01 |
|  | Intercept only | 1 | 166.64 | 11.82 | 0.00 |

**Table 3.** Standardized model-averaged regression coefficients (betas [β]) and 95% confidence limits used to estimate effects of predictors and precision of effects across three candidate model sets. Dashes indicate that a parameter was not present in a final model set. NA values indicate that a parameter was not tested due to absence of certain haemosporidian genera from sites. For all models, Site: Capitan was the reference category.

| Parameter | *Parahaemoproteus* models |  |  | *Plasmodium* models |  |  | *Leucocytozoon* models |  |  |
| --- | --- | --- | --- | --- | --- | --- | --- | --- | --- |
|  | <u>Model-averaged estimate and 95% confidence limits</u> |  |  | <u>Model-averaged estimate and 95% confidence limits</u> |  |  | <u>Model-averaged estimate and 95% confidence limits</u> |  |  |
| | $\beta$ | Lower CL | Upper CL | $\beta$ | Lower CL | Upper CL | $\beta$ | Lower CL | Upper CL |
| Intercept | -0.12 | -1.17 | 0.93 | -2.63 | -3.21 | -2.04 | -1.3 | -2.57 | -0.03 |
| Elevation | -1.13 | -1.93 | -0.33 | - | - | - | 0.77 | 0.02 | 1.53 |
| Latitude | 0.72 | 0.00 | 1.43 | - | - | - | - | - | - |
| Temperature-aridity Index | 1.10 | 0.36 | 1.85 | - | - | - | -0.75 | -1.52 | 0.02 |
| Seasonality Index | 0.77 | -0.42 | 1.96 | - | - | - | - | - | - |
| Site: Chuska | 4.23 | 0.24 | 8.22 | NA | NA | NA | -1.34 | -3.07 | 0.39 |
| Site: Jemez | 4.21 | 0.02 | 8.41 | NA | NA | NA | 0.42 | -1.22 | 2.07 |
| Site: Mt. Taylor | 4.06 | 0.38 | 7.74 | NA | NA | NA | 1.15 | -0.35 | 2.64 |
| Site: Sangre de Cristo | 2.99 | -0.46 | 6.44 | - | - | - | -1.41 | -3.80 | 0.98 |
| Site: San Mateo | 1.79 | -0.39 | 3.97 | - | - | - | 0.05 | -1.65 | 1.74 |
| Site: White | 2.14 | -0.33 | 4.61 | - | - | - | -0.69 | -2.45 | 1.06 |
| Site: Zuni | 5.14 | 1.48 | 8.81 | - | - | - | NA | NA | NA |

### Diversity Analyses

We calculated parasite alpha diversity (within-site haplotype diversity) with rarefaction and the Chao1 index, using EstimateS v9.1.0 (Colwell 2013; Appendix S1). For beta-diversity analyses, we rarefied the data using the R package ‘vegan’ (Oksanen *et al.* 2017) to account for variable sampling. We estimated the haemosporidian phylogeny using maximum-likelihood (ML) in RAxML, v8.2.10 (Stamatakis 2014). We used the GTR+G model of nucleotide substitution and conducted a rapid bootstrap analysis with 1000 replicates, after which we searched for the best-scoring ML tree. We rooted the tree with *Leucocytozoon* (Galen *et al.* 2018). We used the Jaccard index, a presence-absence-based dissimilarity metric (Magurran 2004), to evaluate non-phylogenetic diversity. We used unweighted UniFrac to assess phylogenetic diversity, which measures the fraction of unshared branch lengths between two communities (Lozupone *et al.* 2010).

To characterize host turnover, we used eBird data (eBird.org) and expert knowledge (MJB) to generate bird community lists comprised of the breeding bird species occurring >2,100 m elevation in each sky island (Table S4); we then generated a phylogenetic tree for the sky-island breeding bird community using BirdTree.org, with ‘Hackett All Species’ as the source of trees (see Appendix S1 for details).

### Geographic Structure and Turnover of Parasite Communities

We characterized parasite and bird community composition using nonmetric multidimensional scaling (NMDS), based on unweighted UniFrac distance matrices, in R (Oksanen *et al.* 2017). NMDS was chosen because it uses rank orders to account for large differences in count data and does not assume linear relationships (Winter *et al.* 2017). Communities were more similar when phylogeny was taken into account (Table 4), so we used only unweighted UniFrac distance matrices as inputs for NMDS analyses.

**Table 4.** Measures of non-phylogenetic and phylogenetic turnover among (a) the haemosporidian haplotype community and (b) the breeding bird community. Shaded values (below the diagonals): Jaccard index pairwise beta diversity estimates. Unshaded values (above the diagonals): unweighted UniFrac pairwise beta diversity estimates. Larger numbers indicate greater differences in community structure (i.e. greater turnover).

| Clade | Sky island | Capitan | Chuska | Jemez | Mt. Taylor | Sangre de Cristo | San Mateo | White | Zuni |
| --- | --- | --- | --- | --- | --- | --- | --- | --- | --- |
| **Parasites** | **Capitan** | – | 0.66 | 0.42 | 0.87 | 0.39 | 0.60 | 0.67 | 0.83 |
|  | **Chuska** | 0.90 | – | 0.52 | 0.81 | 0.50 | 0.55 | 0.87 | 0.82 |
|  | **Jemez** | 0.73 | 0.84 | – | 0.79 | 0.07 | 0.52 | 0.82 | 0.80 |
|  | **Mt. Taylor** | 0.84 | 0.81 | 0.74 | – | 0.78 | 0.73 | 0.65 | 0.67 |
|  | **Sangre de Cristo** | 0.57 | 0.89 | 0.69 | 0.82 | – | 0.49 | 0.82 | 0.79 |
|  | **San Mateo** | 0.75 | 0.91 | 0.71 | 0.86 | 0.70 | – | 0.71 | 0.69 |
|  | **White** | 0.75 | 0.86 | 0.78 | 0.81 | 0.70 | 0.71 | – | 0.79 |
|  | **Zuni** | 0.85 | 0.75 | 0.78 | 0.81 | 0.70 | 0.80 | 0.80 | – |
| **Birds** | **Capitan** | – | 0.04 | 0.13 | 0.03 | 0.14 | 0.04 | 0.11 | 0.04 |
|  | **Chuska** | 0.11 | – | 0.11 | 0.02 | 0.12 | 0.08 | 0.09 | 0.03 |
|  | **Jemez** | 0.20 | 0.14 | – | 0.12 | 0.01 | 0.16 | 0.11 | 0.14 |
|  | **Mt. Taylor** | 0.08 | 0.05 | 0.15 | – | 0.13 | 0.07 | 0.10 | 0.02 |
|  | **Sangre de Cristo** | 0.23 | 0.16 | 0.04 | 0.18 | – | 0.17 | 0.11 | 0.15 |
|  | **San Mateo** | 0.07 | 0.15 | 0.23 | 0.12 | 0.26 | – | 0.06 | 0.05 |
|  | **White** | 0.19 | 0.15 | 0.12 | 0.16 | 0.16 | 0.13 | – | 0.09 |
|  | **Zuni** | 0.07 | 0.08 | 0.17 | 0.03 | 0.21 | 0.10 | 0.16 | – |

To quantify the best environmental and geographic predictors of parasite and bird community composition, we used generalized dissimilarity modeling (GDM) in the R package ‘gdm’ (Manion *et al.* 2018). In GDM, predictor variables are transformed using a series of I-spline functions with a high degree of smoothness at places where polynomial pieces connect (“knots”), and models are fit using ML estimation (Ferrier et al., 2007; Glassman *et al.* 2017). The sum of three coefficients for each variable describes the proportion of turnover explained by that variable as determined by the maximum height of its I-spline (Ferrier *et al.* 2007; Fitzpatrick *et al.* 2013). The I-spline slope indicates the rate of turnover along that particular environmental or geographic gradient (Fitzpatrick *et al.* 2013; Glassman *et al.* 2017). The difference in height between any two points along the I-spline corresponds to the modeled contribution of that predictor variable to the difference between those sites; in this way, I-splines indicate the importance of each variable for community composition.

We fit GDMs for the parasite haplotype community and bird community. We started with a full model that included pairwise geographic distance, elevation, temperature-aridity index (PC1), and seasonality index (PC2); for the parasite models we also included the mean values of bird community composition (MDS1 and MDS2); and for bird models we used parasite community composition (MDS1 and MDS2). We used backward elimination to arrive at a best-fit model: beginning with the full model, we removed the variable with the lowest sum of coefficients at each step, and calculated the change in deviance explained (Ferrier *et al.* 2007). For each GDM analysis, results included: (i) a set of best predictor variables, (ii) a fitted I-spline for each predictor variable describing its relationship with turnover, and (iii) percent deviance explained by the model (used to assess GDM model fit).

## RESULTS

### Haemosporidian Abundance and Diversity

We recovered 46 haemosporidian mtDNA haplotypes from 178 birds (Fig. 2; Table S2). A total of 109 birds (61.2%) had at least one haemosporidian infection. We identified 158 infections, and 67% of the mtDNA haplotypes (31 of 46) were novel (Table S5). *Parahaemoproteus* was the most abundant and diverse genus, followed by *Leucocytozoon*; *Plasmodium* was relatively rare. There were no significant differences in haplotype or haplogroup richness among sky islands (Fig. S2).

**Fig. 2.**
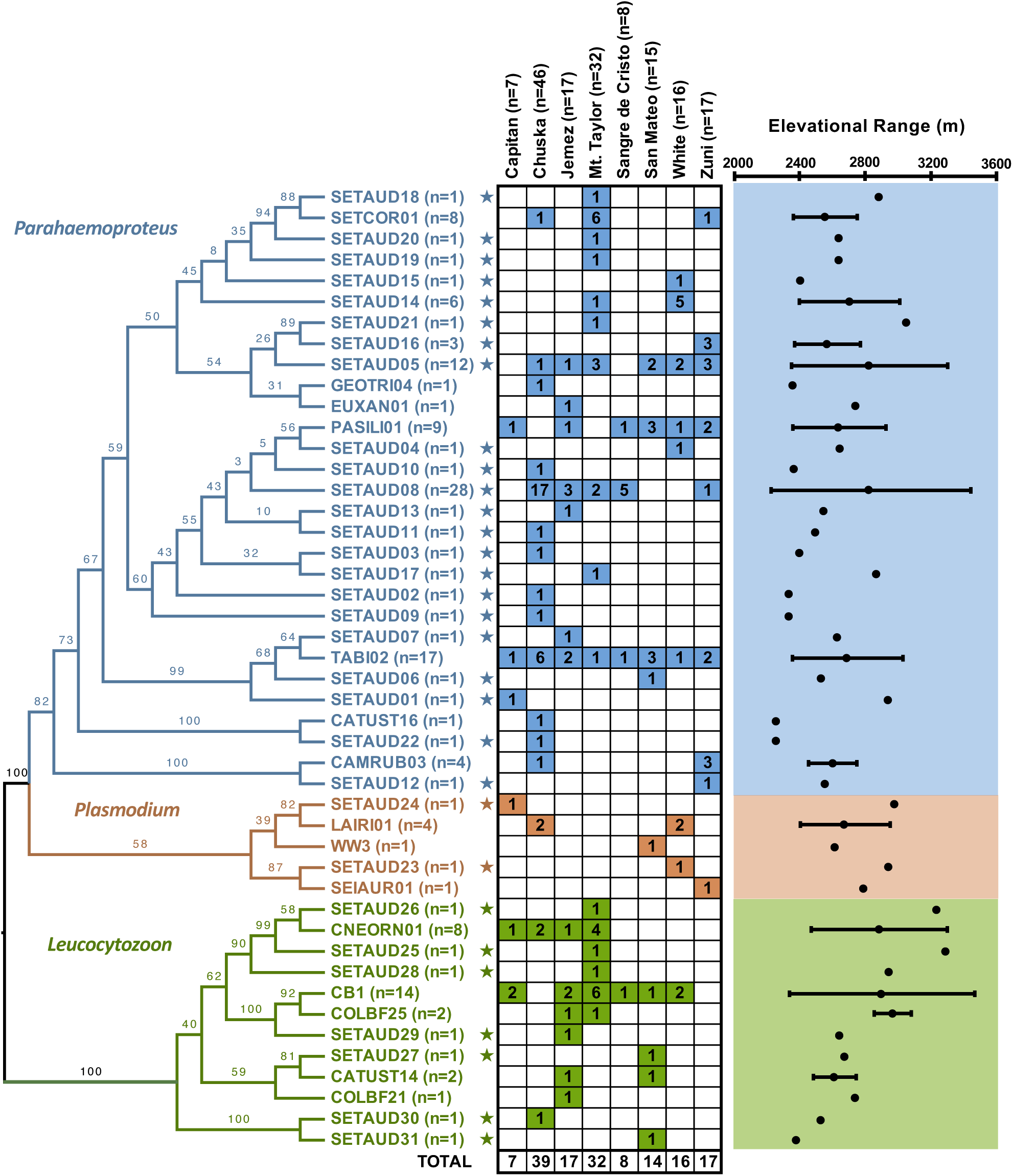
Phylogeny of 46 haemosporidian *cyt b* haplotypes and elevational ranges over which each was encountered in *S. auduboni* from sampled sky islands. Branch labels indicate bootstrap values. Colors correspond to parasite genera. The number of infections is indicated by n, and numbers in the colored cells indicate the number of infections of each haplotype per mountain range. Stars indicate novel haplotypes. The observed elevational range of each haplotype is represented by shaded areas (right); a single dot represents a single elevation.

To account for the possibility that some haplotypes were within-species variants, we collapsed haplotypes into 30 haplogroups. Consolidation reduced diversity by ~1/3, with similar effects on each genus (Fig. S1; Table S3). All results for haplotypes and haplogroups were qualitatively similar, so henceforth we emphasize the haplotype results.

### What Factors Explain Haemosporidian Infection Status?

Predictors of infection status differed among haemosporidian genera. For *Parahaemoproteus* and *Leucocytozoon*, best-fit models performed substantially better than the null, intercept-only model (Table 2). *Parahaemoproteus* infections increased with decreasing elevation and increasing latitude, temperature-aridity index (PC1), and seasonality index (PC2; Tables 2–3). *Plasmodium* infection status was unpredictable, though its relative rarity likely limited statistical power. *Leucocytozoon* infection status varied strongly by mountain range but was not predicted by environmental characteristics (Table 2).

### Host and Parasite Turnover

We found strikingly high rates of parasite turnover among sky islands, far higher than the rates of bird turnover (Table 4). This was consistent for phylogenetic and non-phylogenetic measures of turnover, respectively (Tables 4 and S6). NMDS ordinations confirmed sparse clustering of mountain ranges in parasite ordinations (Figs. S3-S4) and tight clustering of mountain ranges in bird-community ordinations (Fig. S5), consistent with observed high dissimilarity of parasite communities and lower dissimilarity of bird communities (Table 4).

GDM disentangled the effects of geographic and environmental predictors on parasite turnover and bird turnover, respectively (Fig. 3). The GDM model for parasite community turnover explained a moderate proportion of the variance among parasite communities (deviance explained = 37.6%; Table 5). Elevation was the best predictor of parasite community composition, followed by bird community composition, and seasonality index (PC2; Table 5; Figs. 3c and 3e). Although our model included it, pairwise geographic distance between sky islands did not explain any of the variation in parasite community composition.

**Fig. 3.**
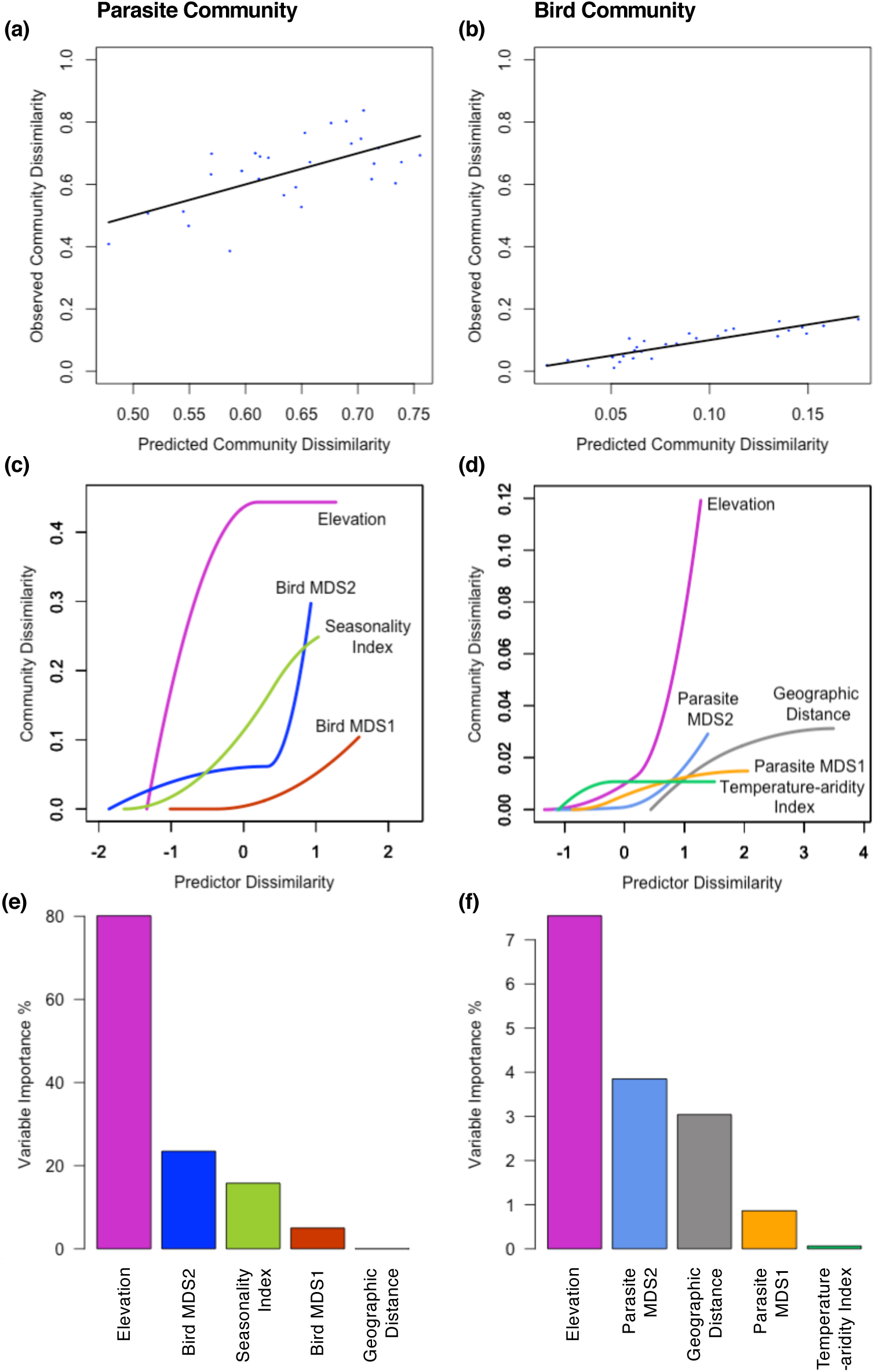
Relationships between observed and predicted community dissimilarity for (a) the parasite haplotype community and (b) the bird community, based on generalized dissimilarity modeling (GDM) analysis. For GDM-fitted I-splines (partial regression fits) for variables associated with (c) parasite beta diversity and (d) bird beta diversity, the maximum height reached by each curve indicates the total amount of community turnover associated with that variable (i.e., its relative contribution to explaining beta diversity). The shape of each I-spline indicates how the rate of community turnover varies with increasing differences in a given predictor variable between sites. Predictor values are taken from x-values of fitted I-splines; these indicate the rate of turnover among mountain ranges for each of the predictors plotted. Variable importance, or percent change in deviance explained by the full model and the deviance explained by a model fit with that variable permuted, is shown for (c) the parasite community model and (d) the bird community model. All predictors depicted are from the best-fit bird and parasite GDM models.

**Table 5.**
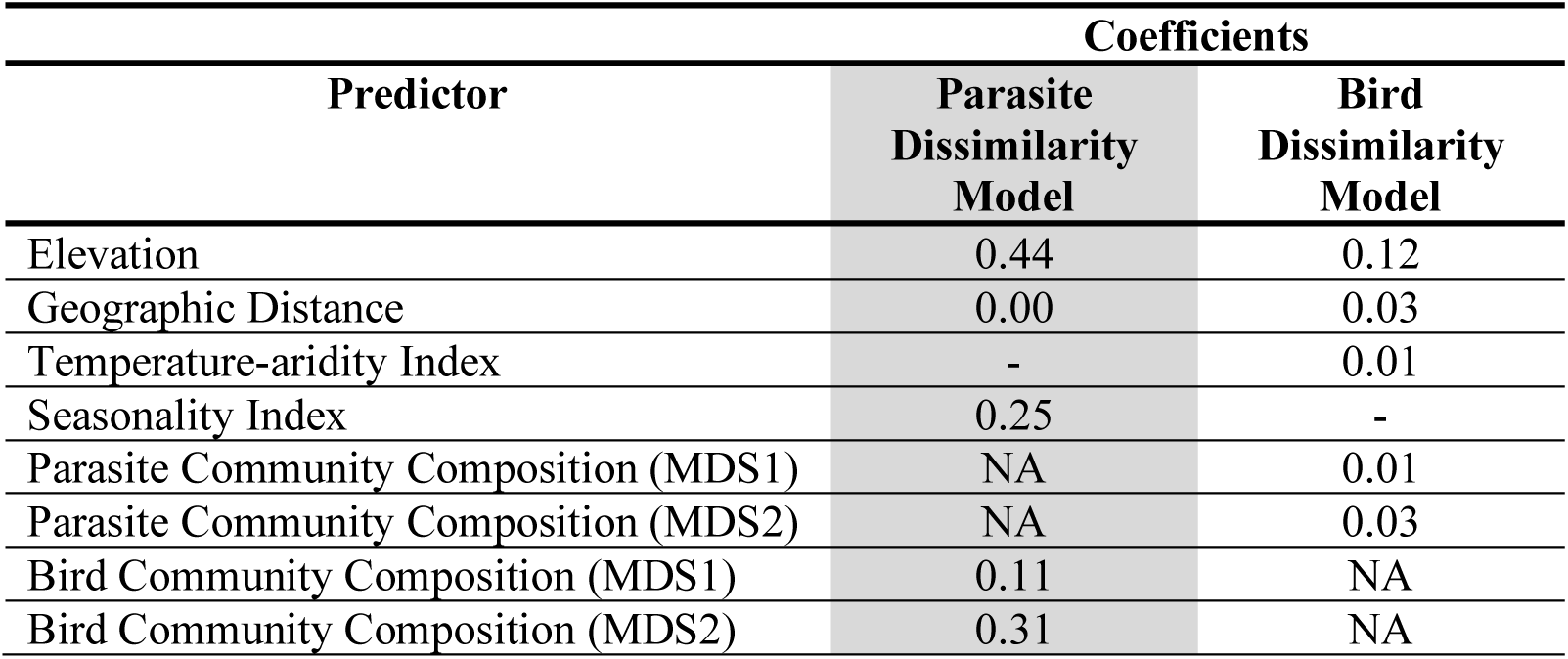
Best models from generalized dissimilarity modeling (GDM) of parasite haplotype (deviance explained = 37.55%) and bird community dissimilarity (deviance explained = 75.61%). Only the predictors and standardized coefficients from the top GDM models for parasite and bird communities are reported. All predictors were standardized prior to input in models. We calculated coefficient values by taking the sum of the three I-spline coefficients, equivalent to the max height on the I-spline curve for each predictor. Dashes indicate that a predictor was not present in a final model. NA values indicate that a predictor was not tested in a model set.

| Predictor | Coefficients |  |
| --- | --- | --- |
|  | Parasite Dissimilarity Model | Bird Dissimilarity Model |
| Elevation | 0.44 | 0.12 |
| Geographic Distance | 0.00 | 0.03 |
| Temperature-aridity Index | - | 0.01 |
| Seasonality Index | 0.25 | - |
| Parasite Community Composition (MDS1) | NA | 0.01 |
| Parasite Community Composition (MDS2) | NA | 0.03 |
| Bird Community Composition (MDS1) | 0.11 | NA |
| Bird Community Composition (MDS2) | 0.31 | NA |

The GDM model of bird community composition explained a large proportion of the variance (deviance explained = 75.6%; Table 5). Model selection by backward elimination retained elevation, parasite community composition (MDS1 and MDS2), geographic distance, and temperature-aridity index (PC1) as the best predictors of bird community composition (Table 5; Fig. 3f).

## DISCUSSION

### Haemosporidian Communities in Sky-Island Audubon’s Warblers

The levels of diversity and novelty in the haemosporidian parasites of sky-island *S. auduboni* were notably high, with 46 total haplotypes recovered, including 31 that were novel. The total number of haemosporidian haplotypes identified in the Yellow-rumped Warbler complex (*S. coronata* and *S. auduboni*) is now 64, making it the world’s second most parasite-rich host, after only *Parus major* (66 haplotypes), according to the MalAvi database. Some of this diversity was attributable to parasite community differences among sky islands, highlighting the need to understand the spatial scale and drivers of parasite species turnover.

The haemosporidian parasite lineages infecting *S. auduboni* included a mix of widespread host generalists and range-restricted host-specialists (Appendix S2). Of the 15 previously published haplotypes we recovered, five spanned multiple continents, and three were found in >15 host species (Table S2). Only one of these widespread and host-generalist lineages was abundant in our survey, CB1 (*Leucocytozoon*), which is known from eight host species, three countries, and two continents (Table S2). The *Plasmodium* haplotypes we identified were already known to be widespread host generalists, consistent with previous studies (Sehgal *et al.* 2010; Table S2). By contrast, four of the 15 previously described haplotypes are known to infect only two host species, and two others are only known from *S. auduboni* (Table S2).

We found that the most abundant parasite lineages in the community appear to be host-specialized. Our two most frequent haplotypes were TABI02 and SETAUD08 (both *Parahaemoproteus*). TABI02 is known only from five species of warblers (Parulidae) in Arizona, Missouri, and New Mexico, whereas SETAUD08 is known only from *S. auduboni*. This is inconsistent with Hellgren, Pérez-tris & Bensch (2009), who showed that host-generalist parasites tended to also be more abundant in any specific host. It is partly consistent with Drovetski *et al.* (2014), who found that the most abundant lineages in Europe included both generalist and specialists. These conflicting results show that the relationship between host-specificity and abundance needs further study.

### Predictors of Haemosporidian Infection Status

The factors that predicted probability of infection differed among haemosporidian genera. *Parahaemoproteus* was the most common and abundant genus encountered in the sky-island forests, consistent with previous findings that it is associated with woodland habitats (Clark *et al.* 2016; Illera *et al.* 2017) and mountains (Lutz *et al.* 2015). Predictors of *Parahaemoproteus* infection status included elevation, seasonality index (PC2), and temperature-aridity index (PC1). The fact that elevation was an important predictor of infection suggests that intensity of parasite pressure varies with elevation, potentially contributing to high host-species turnover along these gradients (Fig. 2). We observed that *Parahaemoproteus* infection increased with increased temperatures and decreased precipitation (temperature-aridity index; Table 3), although our study was restricted montane woodlands that are relatively cool and wet. Zamora-Vilchis *et al.* (2012) also found that *Parahaemoproteus* was strongly positively correlated to temperature in the Australian Wet Tropics bioregion, where mean annual temperature of sampled localities ranged from 16.4–21.8°C. Illera *et al.* (2017) found that temperature was negatively correlated with *Parahaemoproteus* prevalence in one mountain range in northern Spain, where mean annual temperature was ~15°C. Recently, Pulgarín-R *et al.* (2018) found no association between *Parahaemoproteus* prevalence and climatic variables along a tropical aridity gradient in Colombia.

We found that *Plasmodium* infection status was unpredictable, likely due to its low frequency in our survey. The relative rarity of *Plasmodium* is consistent with previous studies in the southwestern United States (Marroquin-Flores *et al.* 2017) and other arid environments (Zamora-Vilchis *et al.* 2012).

*Leucocytozoon* varied idiosyncratically by mountain range (Table 2). Although environmental variables were not included in top models, there was a trend toward *Leucocytozoon* infection increasing at higher elevations and with cooler, wetter conditions (i.e. temperature-aridity index; Table 3). These effects may have been modest because the entirety of our survey took place in relatively cool and forested montane habitats, but they are consistent with studies from across the globe: Lutz *et al.* (2015) found that rates of *Leucocytozoon* infection decreased at drier, low elevation sites in Malawi; Galen & Witt (2014) found more *Leucocytozoon* in wetter conditions and at higher elevations in Peru; and Merino *et al.* (2008) found that *Leucocytozoon* prevalence increased with latitude in Chile. It is also likely that *Leucocytozoon* variation among sky-island mountain ranges is partly attributable to un-modeled aspects of the environment, such as proximity to water, which could affect local abundance of simulid vectors.

### Patterns and Predictors of Haemosporidian Parasite Turnover

We found that parasite communities exhibited striking turnover, far greater than that of bird communities (Fig. 3a-b; Table 4). Previous studies have examined the scale at which haemosporidian parasite community turnover occurs, but at much larger spatial scales, ranging from ~750 km (Ellis *et al.* 2015) to ~2,400 km (Scordato & Kardish 2014). Low turnover has been demonstrated at spatial scales as large as ~2110 km (Fecchio *et al.* 2017b). Here we describe turnover of host and parasite communities at relatively fine spatial scales of a few hundred kilometers. The fact that haemosporidian parasite turnover is high at this fine scale, across which avian host communities are relatively constant, suggests that haemosporidian communities exhibit turnover patterns closer to the ‘micro-scale’ expected of microbial communities (Astorga *et al.* 2012; Nemergut *et al.* 2013). Rather than parasite turnover mirroring host turnover, it was far greater (Fig. 3a-b; Table 4). This is consistent with fundamentally different processes causing turnover for micro- (parasite) and macro-organisms (hosts). For example, parasite communities might turn over faster than host communities due to higher sensitivity to climate variation, faster evolutionary rates, or host-switch driven speciation. The finer spatial scale of turnover in parasites could exert spatially varying selective pressure on host immune systems, promoting host genetic diversity and potentially accelerating host diversification (Laine & Tellier 2008; Thornhill & Fincher 2013; Betts *et al.* 2018).

We found that parasite turnover is predicted by host turnover. This finding, consistent with previous studies that encompassed equivalent and/or larger spatial scales (Ishtiaq *et al.* 2010; Svensson-Coelho & Ricklefs 2011; Ellis *et al.* 2015; Clark & Clegg 2017; Fecchio *et al.* 2017b) suggests that at least some of the geographic variation in host communities is a cause or consequence of heterogeneity of parasite communities. It further implies that the identity of a specific species in a multi-host, multi-parasite system is important for the persistence (or lack of persistence) of its potential symbiont species. This is consistent with indirect antagonistic effects among bird or parasite species, respectively, mediated by symbiont species. Such effects could limit ranges and ultimately accelerate diversification for hosts or parasites.

Although differences among sky-island parasite communities were somewhat idiosyncratic, turnover was partly explained by abiotic environmental variables and biotic ecological variables (Fig. 3). Specifically, parasite turnover was best explained by the combination of elevation, seasonality index (PC2), and bird community composition (bird MDS1 and bird MDS2; Fig. 3a). Although a distance-decay relationship is expected to exist among parasite communities at large spatial scales (Nekola & White 1999; Poulin 2003), geographic distance did not explain parasite turnover among sky islands. This negative finding is consistent with Fecchio et al. (2017b), who found no effect of geographic distance on haemosporidian community turnover in manakins across the Amazon basin; however, Fecchio *et al.* (2017a) did find a latitude effect on turnover among Amazonian areas of endemism in a broader community survey. The lack of a distance effect at this scale suggests that dispersal-distance (and colonization potential) of parasites does not constrain the island-biogeographic cycle by which sky-island communities are assembled; rather, environmental filters are dominant drivers of parasite community composition. This fits with the idea that vagile, migratory hosts such as *S. auduboni* provide long-distance dispersal capability for parasites, even if the potential is seldom realized due to ecological constraints on colonization.

In our study, we sampled haemosporidian parasites from a single, widespread host. Given the tendency of haemosporidians to infect multiple host species (Galen & Witt 2014; Moens *et al.* 2016), and given that *S. auduboni* shares many parasite haplotypes with other New Mexico breeding bird species (Marroquin-Flores *et al.* 2017), we expect that the patterns observed in *S. auduboni* reflect those of the whole bird community. This rationale is supported by Svensson-Coelho & Ricklefs (2011), who found that a single host species, the Black-faced Grassquit, reflected diversity and turnover patterns in the whole haemosporidian community. The high parasite turnover we observed within a single host species might remain the same or become greater if the entire breeding bird community were sampled. On the other hand, variation in Northern Cardinal haemosporidians from place to place (Ellis *et al.* 2015) was greater than expected based on a variation among a suite of sampled host species, suggesting that a single host species approach could result in inflated measures of turnover. Future studies should examine whole bird communities, but there are few such surveys at present, particularly in continental systems, because of the sampling efforts required.

Our application of GDM yielded new insights regarding predictors of community turnover (Fig. 3; Table 5), and this method seems ideally suited for the further study of avian haemosporidians. The advantages of using GDM included adequately accounting for geographic distance, curvilinear effects, and collinearity among predictors. By simultaneously accounting for environment, host and parasite community composition, and expected non-linear relationships of community dissimilarity with environmental and geographic distance, our GDM results explained substantial variation in parasite (38%) and bird communities (76%), respectively.

### Conclusions

This study revealed causes of abundance and turnover for haemosporidian parasite communities across an array of sky islands in southwestern North America. Infection status was predicted by environmental characteristics, but sky islands also showed idiosyncratic variation. Parasite turnover was, in nearly all cases, three-fold higher than bird turnover at spatial scales of only a few hundred kilometers. Generalized dissimilarity modeling (GDM) revealed that variation among sky-island parasite community composition could be explained by a combination of abiotic and biotic ecological variables, but not by geographic distance. Importantly, we found that parasite turnover and host turnover are linked; this finding was surprising, though not unprecedented, because both parasites and hosts tend to be generalized in these communities. This implies that the identities of specific host and parasite species matter to community composition, even in a complex multi-host, multi-parasite system.

## ACKNOWLEDGEMENTS

We thank Andrea N. Chavez, Zac A. Cheviron, Ariel M. Gaffney, Andrew B. Johnson, Rosario Marroquin-Flores, Laura Pagès Barceló, Rachel Price, Paulo Pulgarín, Jennifer Rudgers, C. Gregory Schmitt, and Ashley Smiley. This research was supported by the National Science Foundation (NSF DEB-1146491), the Federal Bureau of Land Management Rio Puerco Field Office, the New Mexico Ornithological Society. Collecting permits were granted by the U.S. Fish and Wildlife Service (MB-726595-0), New Mexico Department of Game and Fish (Auth. No. 3217), Arizona Department of Game and Fish (SP612810), and Navajo Nation Department of Fish and Wildlife (716-2012 and 663-2013).

## DATA ACCESSIBILITY

Specimen and parasite data are available as Supplemental Files. Parasite sequences are archived in ARCTOS, MalAvi, and GenBank (Accession numbers MF752555–MF752704).

## AUTHOR CONTRIBUTIONS

CCW, CJW, and JLW conceived and designed the study; CJW, CCW, CJS, DCS, MBJ, LNB, SCG, and JLW collected data; JLW, CCW, LNB, and ASW analyzed the data; JLW, CCW, and LNB wrote the manuscript with input from all authors.

